# Different forms of fear extinction are supported by distinct cortical substrates

**DOI:** 10.1101/2020.01.27.921221

**Authors:** Belinda P. P. Lay, Audrey Pitaru, Nathan Boulianne, Guillem R. Esber, Mihaela D. Iordanova

**Affiliations:** Center for Studies in Behavioural Neurobiology, Department of Psychology, Concordia University, Montreal, QC, Canada; Department of Psychology, Brooklyn College of the City University of New York, Brooklyn NY, USA

## Abstract

Understanding how learned fear can be reduced is at the heart of treatments for anxiety disorders. Tremendous progress has been made in this regard through extinction training in which an expected aversive outcome is omitted. However, current progress almost entirely rests on this single paradigm, resulting in a very specialized knowledgebase at the behavioural and neural level of analysis. Here, we used a paradigm-independent approach to show that different methods that lead to reduction in learned fear are dissociated in the cortex. We report that the infralimbic cortex has a very specific role in fear reduction that depends on the omission of aversive events but not on overexpectation. The orbitofrontal cortex, a structure generally overlooked in fear, is critical for downregulating fear when fear is inflated or overexpected, but not when an aversive event is omitted.

---

Extinction learning has captivated behavioural and neural science for more than a century. It has done so because it allows for the reduction of behaviours that were once adaptive but are no longer so, and gives the therapist a handle to combat others that were never adaptive in the first place. The most-widely used method for supressing unwanted behaviour relies on the *omission* of the event that drives this behaviour, that is, extinction driven by outcome omission. In the context of fear learning, extinction by omission involves the dramatic reduction in fear-related behaviours typically observed after presenting a previously established signal for an aversive event (i.e., a tone paired with shock; tone→shock) in the absence of that event (tone presented alone; tone→nothing). Given its simplicity and effectiveness in the treatment of anxiety disorders^1–6^, extinction by omission has received significant attention in a quest to understand its underlying behavioural and neural mechanisms^7–15^. Critically, although much progress has been made, this progress is limited to the case of outcome omission, while another equally relevant form of extinction learning that also drives reduction in unwanted behaviour, namely overexpectation, remains largely unexplored. This single-paradigm approach is restrictive because at best it can oversimplify and at worst even misrepresent the function of brain areas implicated in extinction learning. Here, we move beyond this paradigm-specific approach and embarked on an investigation into how the brain learns from extinction using two behavioural designs: extinction driven by outcome omission (described above) and extinction driven by overexpectation (described below).

In overexpectation, reduction in previously established fear responses ensue, strikingly, despite continued delivery of the aversive event. This is possible because separately established signals of a common aversive event (i.e., tone→shock; light→shock) can summate their fear-inducing properties when encountered simultaneously (tone+light), triggering a state of exacerbated fear. The presentation of the same (i.e., unintensified) aversive event in that state (tone+light→shock) engages a self-regulatory mechanism that lessens the exacerbated fear by partially extinguishing the fear elicited by each individual signal^16–18^. Since signals for threat often co-occur (think of the sight of a mic and that of a staring crowd for a glossophobic), failure to reduce fear by overexpectation is a likely contributor to the aberrant and persistent fear characterizing anxiety disorders such as panic disorder, social anxiety disorder or post-traumatic stress disorder.

The infralimbic cortex (IL) is considered to be the key brain locus of extinction learning in fear^8,9,19–24^. Importantly, evidence for this role is exclusively derived from extinction by omission designs. If the IL is critical for learning to reduce fear in general, then it should do so irrespective of the conditions that generate this reduction. That is, the role of the IL should not be limited to the conditions specified by outcome omission, rather it should also be seen in overexpectation. To date, there is no evidence for this generality of IL function. Alternatively, the IL may have a very specific role in learning that leads to suppression in established behaviour when expected outcomes are omitted. To determine if the IL has a paradigm-independent or a paradigm-specific role in fear extinction, we first examined whether the IL is necessary for reduction in fear driven by overexpectation and then by outcome omission.

We implanted sixty-three Sprague Dawley rats with cannula targeting the IL bilaterally (Figures 1A and 1B). Following a week of recovery, the rats were trained to associate a tone with shock (tone→shock) and a flashing light with shock (light→shock, Figure 1C, online methods). Percent time spent freezing to the cues was taken as a measure of the level of conditioned fear^25,26^. Following robust fear conditioning to the individual cues (Figure 1D, see figure legend for statistics), the tone and the light were presented in compound and paired with the same single shock as that delivered during initial fear acquisition in order to generate the overexpectation condition (Figure 1C). Prior to overexpectation training the rats received either infusions of muscimol and baclofen (0.1mM muscimol-1 mM baclofen) into the IL or vehicle. This was done in order to inactive the IL cortex during overexpectation learning and compare its effect to an identical group that received overexpectation training but in the presence of a functional IL. Rats in the control condition did not receive overexpectation training but received identical infusions of the drug or the vehicle (Figure 1C). There were no differences in conditioned fear between the groups during the overexpectation training phase (Figure 1 E, see figure legend for statistics). Following this training, rats received probe testing in which the target cue (tone or light, counterbalanced) was presented alone in the absence of shock (Figure 1C). Fear to the target cue was lower in the overexpectation groups compared to the control groups irrespective of whether IL function was intact during learning (Figure 1E, see figure legend for statistics).

**Figure 1.**
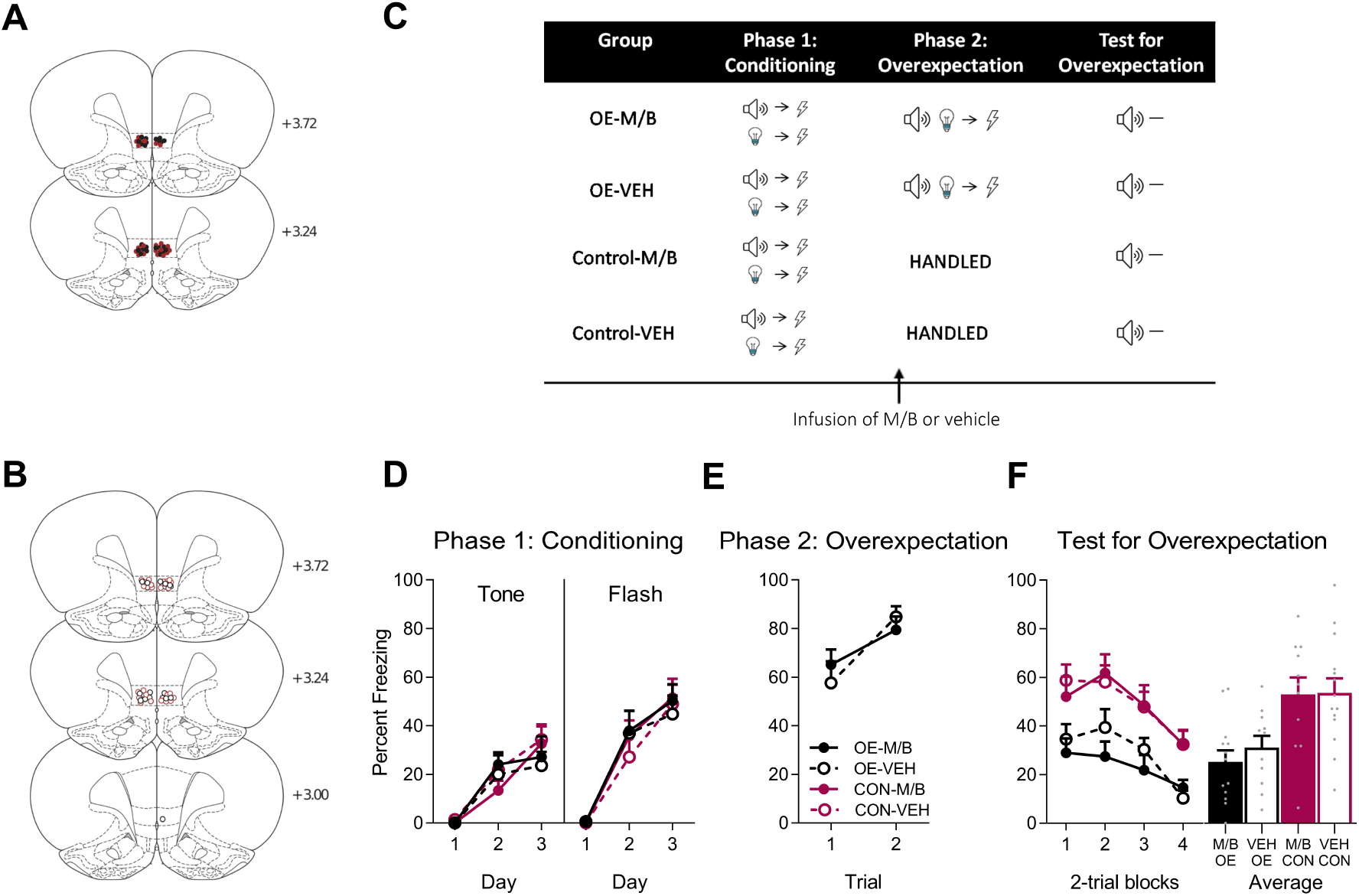
The IL is not necessary for overexpectation. Location of cannula placements for A) drug- and B) vehicle-treated rats in the IL cortex overexpectation experiment as verified based on the atlas of Paxinos and Watson.^65^ The symbols represent the most ventral point of the cannula track for each rat and distances are indicated in millimeters from bregma. Overexpectation-M/B rats are shown with filled black circles, overexpectation-vehicle rats are shown with open circles, control-M/B rats are shown with filled burgundy circles, and control-vehicle rats are shown with open burgundy circles. C) Behavioural protocol for Overexpectation. Rats are trained to associate two individual cues (tone and light, counterbalanced) with a foot-shock US during Phase 1. Immediately prior to overexpectation training (Phase 2), the lateral OFC was pharmacologically silenced with muscimol and baclofen (M/B). Rats in the overexpectation group (OE) received compound presentations of the two cues followed by the delivery of a foot-shock. Rats in the control group (Control) were handled. All rats were then tested for conditioning responding the following day to either tone or light (counterbalanced). Behavioural data are represented as mean +SEM percent levels of freezing. D) Acquisition of conditioned freezing responses to the tone and light was successful (max *F*_(1,45)_ = 166.96, *p* < 0.01, 95% CI [1.79, 2.46], mixed ANOVA) and equivalent across all four groups (max *F*_(1, 45)_ = 0.069, *p* = 0.84, mixed ANOVA) during Phase 1. E) Infusions of M/B in the IL cortex had no effect on within-session responding to the compound presentations across Phase 2 (*F*_(1, 22)_ = 0.022, *p* = 0.88, mixed ANOVA). F) Learning from overexpectation was successful (*F*_(1, 45)_ = 19.21, *p* < 0.001, 95% CI [−1.48, −0.41], d = 1.30, mixed ANOVA). However, there was no effect of IL inactivation on retrieval of the overexpectation memory when tested drug-free (*F*_(1, 45)_ = 0.33, *p* = 0.57, mixed ANOVA).

Following the overexpectation part of the experiment, the same rats took part in an extinction by omission experiment (Figure 2C). The allocation of rats to groups was counterbalanced based on their prior experience in the overexpectation study (see methods) and their placements in this part of the experiment are depicted as per their new group assignment (Figure 2A and 2B). All rats were conditioned to fear a novel cue (steady light or white-noise, counterbalanced, Figure 2D, see figure legend for statistics). This training was followed by extinction training in which the cue was presented in the absence of the associated shock in half of the cohort or no training in the other half (Figure 2E). Again, prior to extinction by omission training, the rats received either inactivating infusions of the IL with muscimol and baclofen or vehicle. Identical infusions were also given to the control rats that did not receive extinction. Inactivation of the IL prior to extinction by omission training slowed down the reduction in freezing compared to the vehicle group (Figure 2E, see figure legend for statistics). The following day, the rats received probe testing in which the cue was presented in the absence of shock. Fear to this cue was lower in rats that received extinction by omission training in the presence of a functional IL compared to rats that received extinction by omission training while the IL was inactivated or rats that received no extinction training at all (Figure 2F). These data provide key evidence for the dissociable role of the IL in fear reduction under conditions of expected shock omission but not of shock overexpectation. That is, the IL is critical to downregulating conditioned fear when threatening events are no longer present, leaving the question of how the cortex downregulates fear under overexpectation unanswered.

**Figure 2.**
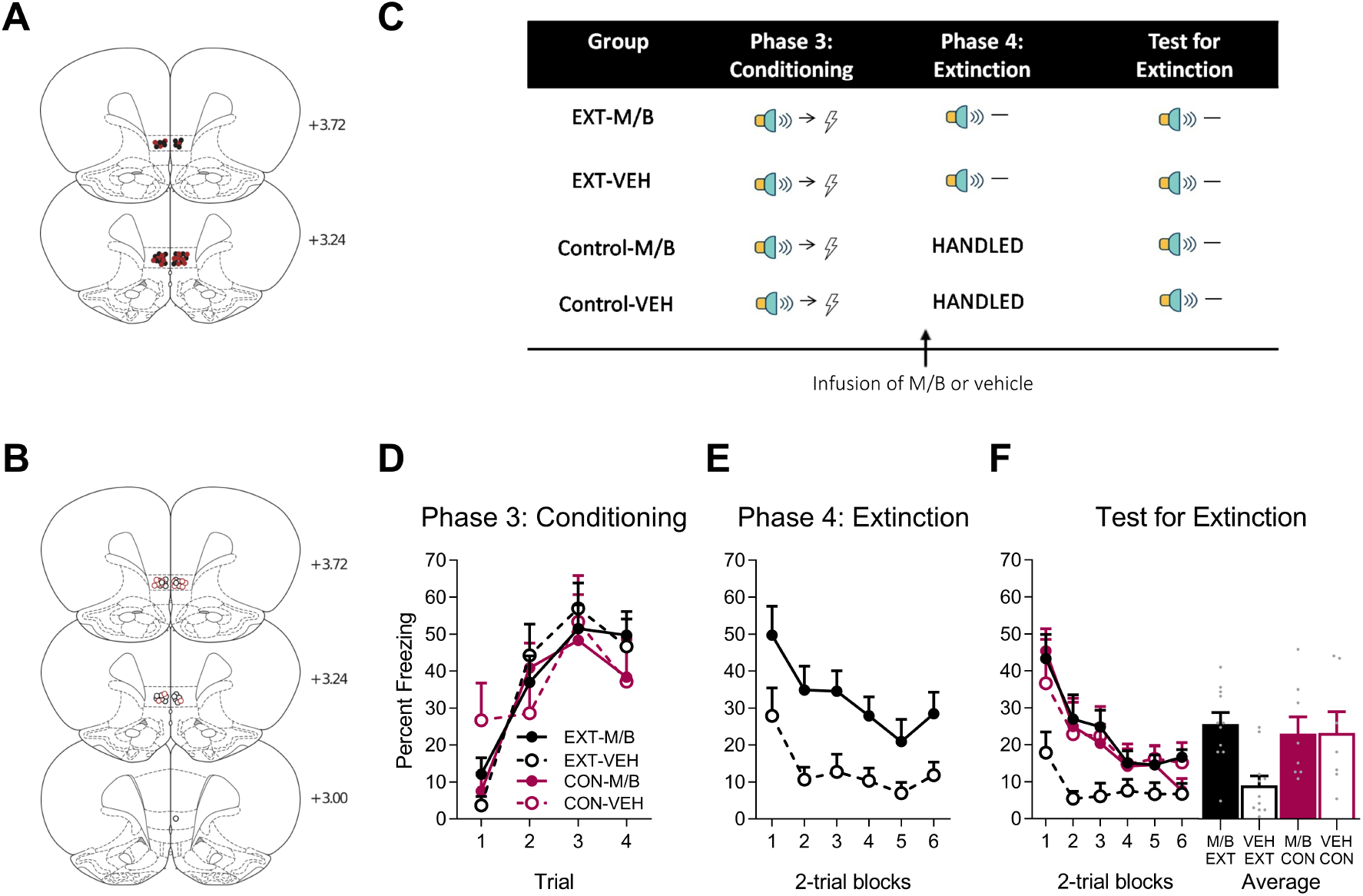
The IL is necessary for extinction by omission. Using the same animals, the location of cannula placements reallocated for A) drug- and B) vehicle-treated rats in the lateral OFC extinction experiment as verified based on the atlas of Paxinos and Watson.^65^ The symbols represent the most ventral point of the cannula track for each rat and distances are indicated in millimeters from bregma. Extinction-M/B rats are shown with filled black circles, extinction-vehicle rats are shown with open circles, control-M/B rats are shown with filled burgundy circles, and control-vehicle rats are shown with open burgundy circles. C) Behavioural protocol for Extinction. Following the overexpectation experiment, rats received conditioning to a novel stimulus (auditory or visual cue, counterbalanced) paired with foot-shock during Phase 3. Prior to extinction in Phase 4, rats received an intra-lateral OFC infusion of M/B or vehicle. Rats in the extinction condition received non-reinforced presentations of the previously fear conditioned stimulus. Rats in the control conditioned were handled. All rats were then tested for conditioned freezing responding the following day. Behavioural data are represented as mean +SEM percent levels of freezing. D) Acquisition of conditioned freezing responses to the novel stimulus across Phase 3 was successful (*F*_(1, 33)_ = 44.82, *p* < 0.01, 95% CI [0.76, 1.42], mixed ANOVA) and equivalent across all four groups (*F*_(1, 33)_ = 0.84, *p* = 0.37, mixed ANOVA). E) Rats infused with M/B in the IL cortex prior to extinction training froze significantly more to the cue compared to vehicle-treated rats (*F*_(1, 20)_ = 12.80, *p* = 0.002, 95% CI [0.40, 1.54], d = 1.53, mixed ANOVA). F) Inactivation of the IL cortex prior to extinction training disrupted retrieval of the extinction memory on drug-free Test: rats treated with M/B during extinction froze significantly more during presentations of the CS at Test compared to rats treated with vehicle (*F*_(1, 33)_ = 11.43, *p* = 0.002, 95% CI [0.16, 1.54], d = 1.85, mixed ANOVA).

To answer this question, we moved beyond the traditional fear circuit. Our candidate was the lateral orbitofrontal cortex (lOFC), a structure strongly linked to reward learning, including the computation of reward value estimates^27^. We targeted the OFC for three reasons. Firstly, it has dense reciprocal projections with the basolateral amygdala (BLA)^32,33^, which has been extensively implicated in fear acquisition and extinction by omission^15^. Secondly, the lOFC has been linked to fear^34–39^ and anxiety^40,41^. Finally, it supports learning from overexpectation of reward^42–44^.

To examine the role of the lOFC in reduction in learned fear responses that ensue from extinction training, rats received identical training to that described above, that is overexpectation followed by extinction by omission, with the exception that they were implanted with bilateral canulae into the lOFC (Figures 3A and 3B). Following fear conditioning with the individual cues (tone→shock, light→shock, Figure 3C and 3D, see legend for statistics), rats were infusions with muscimol and baclofen or vehicle into the lOFC and then trained in overexpectation (tone+light→shock; Figure 3C). Rats in the control conditions received identical infusions but no overexpectation training (see supplementary methods). Freezing to the compound cues during the overexpectation phase was similar between the groups (Figure 3E, see legend for statistics). To determine whether the lOFC was important for learning during overexpectation of fear, rats received a probe test with the target cue (tone or light, counterbalanced between rats) alone in the absence of shock (Figure 3C). Fear to the target cue was lower in rats that received overexpectation training in the presence of a functional lOFC compared to rats that received identical training but in the absence of a functional lOFC or rats that received no overexpectation training at all (Figure 3F, see legend for statistics). That is, in the absence of the lOFC, rats are not able to learn to inhibit conditioned fear responding that normally ensue from overexpectation training.

**Figure 3.**
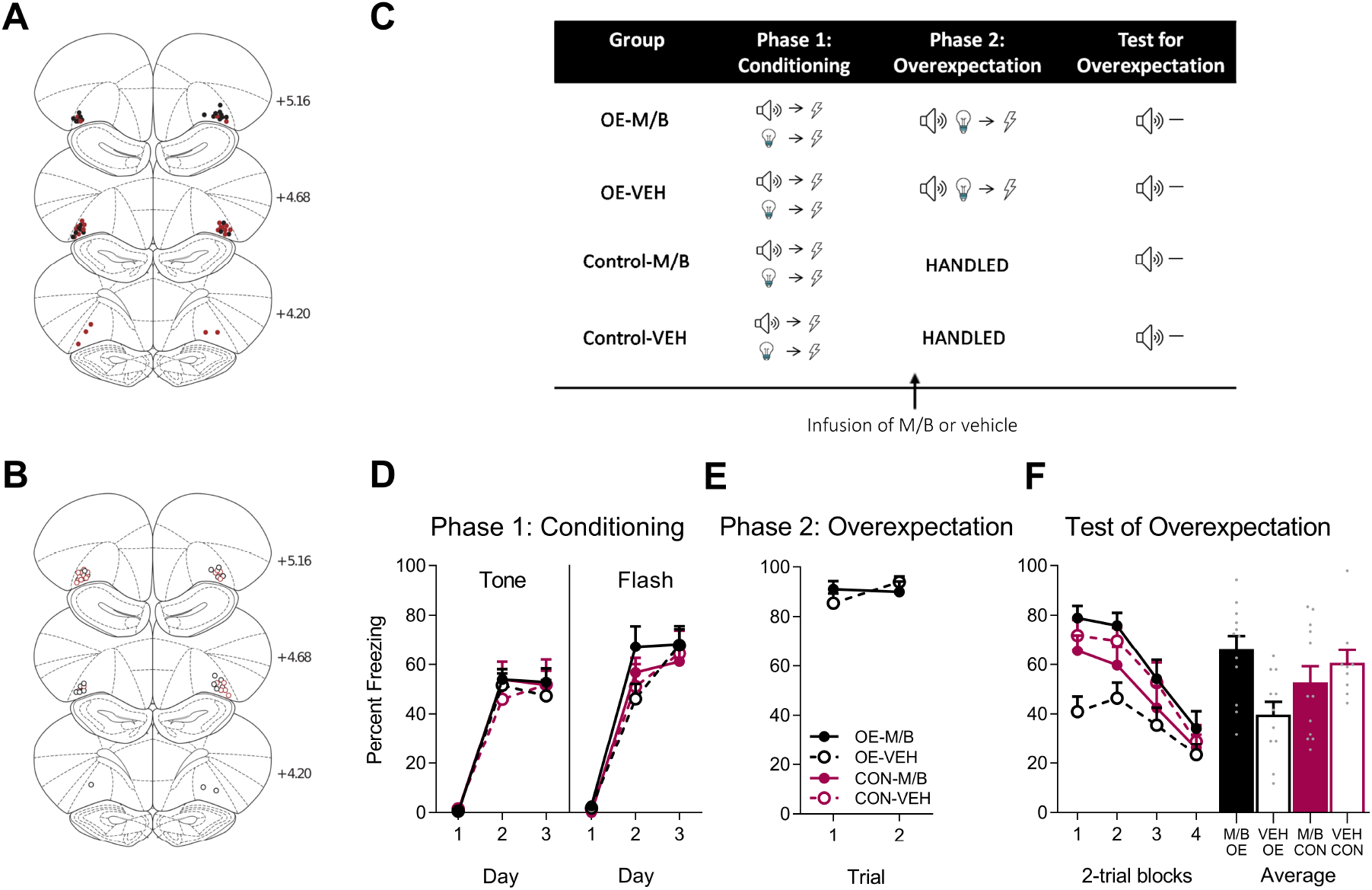
The OFC is necessary for overexpectation. Location of cannula placements for A) drug- and B) vehicle-treated rats in the lateral OFC overexpectation experiment as verified based on the atlas of Paxinos and Watson.^65^ The symbols represent the most ventral point of the cannula track for each rat and distances are indicated in millimeters from bregma. Overexpectation-M/B rats are shown with filled black circles, overexpectation-vehicle rats are shown with open circles, control-M/B rats are shown with filled burgundy circles, and control-vehicle rats are shown with open burgundy circles. C) Behavioural protocol for Overexpectation. Rats are trained to associate two individual cues (tone and light, counterbalanced) with a foot-shock US during Phase 1. Immediately prior to overexpectation training (Phase 2), the lateral OFC was pharmacologically silenced with muscimol/baclofen (M/B). Rats in the overexpectation group (OE) received compound presentations of the two cues followed by the delivery of a foot-shock. Rats in the control group (Control) were handled. All rats were then tested for conditioning responding the following day to either tone or light (counterbalanced). Behavioural data are represented as mean +SEM percent levels of freezing. D) Fear acquisition to the tone and light across Phase 1 was successful (max *F*_(1, 40)_ = 264.90, *p* < 0.01, mixed ANOVA) and equivalent across all four groups (max *F*_(1, 40_ = 0.15, *p* = 0.70, mixed ANOVA). E) Infusions of M/B in the OFC had no effect on within-session responding to the compound stimulus during Phase 2 (*F*_(1, 21)_ = 0.05, *p* = 0.83, mixed ANOVA). F) Inactivation of the lateral OFC prior to overexpectation training disrupted the overexpectation effect on drug-free Test: rats in the overexpectation condition treated with M/B froze significantly more during presentations of the target stimulus at Test compared to vehicle-treated rats (*F*_(1, 40)_ = 10.93, *p* = 0.002, 95% CI [0.18, 1.80], d = 1.44, mixed ANOVA).

Similar to the experiment reported above, the same rats took part in an extinction by omission experiment. The allocation of rats to groups was counterbalanced based on their prior experience in the overexpectation study (see supplemental methods) and their placements in this part of the experiment are depicted as per their new group assignment (Figure 4A and 4B). All rats were conditioned to fear a novel cue (steady light or white-noise, counterbalanced, Figure 4D, see figure legend for statistics). Following this training, half of the cohort of rats received infusions of either muscimol and baclofen or vehicle into the lOFC prior to extinction by omission training in which the cue was presented in the absence of the associated shock (Figure 4C). The rest received identical infusions in the absence of extinction by omission training. Freezing during extinction declined similarly in both groups (Figure 4E, see legend for statistics). The following day, rats received probe testing in which the cue was presented in the absence of shock. Fear to this cue was lower in the rats that received extinction by omission training compared to the controls (Figure 4F) irrespective of whether extinction training took place under a functional or an inactivated lOFC. These data provide clear evidence for a role of lOFC in fear reduction driven by overexpectation but not by outcome omission.

**Figure 4.**
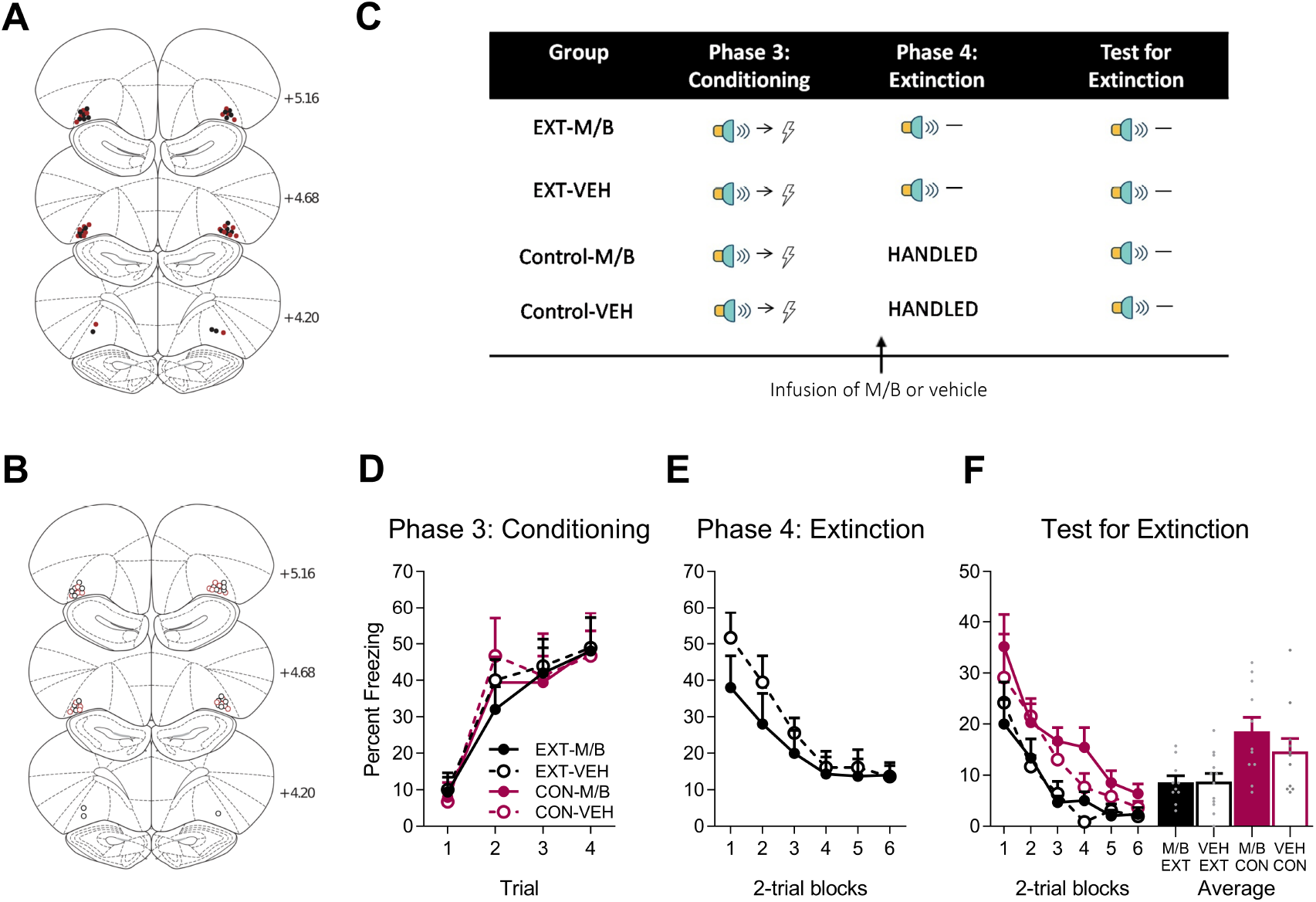
The OFC is not necessary for extinction by omission. Using the same animals, the location of cannula placements reallocated for A) drug- and B) vehicle-treated rats in the lateral OFC extinction experiment as verified based on the atlas of Paxinos and Watson.^65^ The symbols represent the most ventral point of the cannula track for each rat and distances are indicated in millimeters from bregma. Extinction-M/B rats are shown with filled black circles, extinction-vehicle rats are shown with open circles, control-M/B rats are shown with filled burgundy circles, and control-vehicle rats are shown with open burgundy circles. C) Behavioural protocol for Extinction. Following the overexpectation experiment, rats received conditioning to a novel stimulus (auditory or visual cue, counterbalanced) paired with foot-shock during Phase 3. Prior to extinction in Phase 4, rats received an intra-lateral OFC infusion of M/B or vehicle. Rats in the extinction condition received non-reinforced presentations of the previously fear conditioned stimulus. Rats in the control conditioned were handled. All rats were then tested for conditioned freezing responding the following day. Behavioural data are represented as mean +SEM percent levels of freezing. D) Acquisition of conditioned freezing responses to the novel stimulus across Phase 3 was successful (*F*_(1, 40)_ = 67.52, *p* < 0.01, 95% CI [0.93, 1.53], mixed ANOVA) and equivalent across all four groups (*F*_(1, 40)_ = 0.001, *p* = 0.98, mixed ANOVA). E) Infusion of M/B in the lateral OFC prior to extinction training had no effect on within-session performance (*F*_(1, 20)_ = 0.90, *p* = 0.35, mixed ANOVA). F) Extinction training reduced fear on drug-free Test (*F*_(1, 40)_ = 13.65, *p* = 0.001, 95% CI [−0.88, −0.14], d = 1.12, mixed ANOVA) but this extinction effect was not modulated by inactivation of the lateral OFC during extinction learning (*F*_(1, 40)_ = 0.79, *p* = 0.38, mixed ANOVA).

Our results represent a fundamental contribution to the study of fear extinction. By using two different types of extinction, we show that the brain downregulates fear in a paradigm-specific manner. This is important because it flies in the face of a parsimonious explanation for extinction in the IL and lOFC. Specifically, both extinction by omission and by overexpectation are underscored by the same negative prediction error mechanism^45^, that is when expectations surpass reality. While this common process can account for the behavioural effect of reduced fear on test, is unlikely to be what is regulated by the IL and OFC or we would have seen similar disruptions in both behavioural designs. In other words, the neural dissociation of these tasks suggests that the lOFC and the IL support extinction learning via different task-dependent processes. Therefore, it is important to underscore that the discrepancy between real and expected events is generated differently in the two designs. Our data show that the IL is important for reduction in learned fear when the expectation is held constant but the delivery of the aversive event is manipulated (through omission), whereas the lOFC is critical for fear reduction when the delivery of the aversive event is held constant but the expectation is manipulated (through overexpectation). This is supported by studies in the appetitive field: Disruption of IL function results in deficits in extinction of appetitive conditioning,^46–49^ and extinction of drug-seeking,^50,51^ whereas disruption of OFC function leads to deficits in appetitive overexpectation.^42–44^

The specific involvement of the IL in reduction of learned fear through extinction by omission but not overexpectation provides important insight into the type of inhibitory process that the IL may regulate. One possibility is that in extinction by omission the reduction in the aversive state of the animal as a result of outcome (shock) omission leads to the development of an inhibitory association between the cue and the state of fear. In overexpectation, such an inhibitory association is unlikely as the aversive outcome continues to be delivered, thus encouraging the development of inhibitory associations between the cue and the second expected but not delivered aversive event. This speculation would require further examination but a similar argument has been made in the appetitive context^49^. This account would also suggest that the differential role of the lOFC in overexpectation but not extinction by omission could be due to the development of inhibitory cue-outcome associations but not inhibitory associations between the cue and a generalized state of fear. Anatomical evidence is somewhat in line with this idea. The IL innervates the central amygdala via the intercalated cells masses,^52^ a structure heavily implicated in gating the expression of conditioned fear responses^53–57^. The lOFC has dense reciprocal projections with the BLA^32,33^ which are important for representing cue-outcome associations^58,59^ and distinct BLA neurons show elevated neural firing to cues that have undergone extinction training.^15^ Alternatively, the lOFC may be critical for the flexible integration of associations that lead to enhanced fear in overexpectation which is key to generating a negative prediction error and driving a reduction in conditioned responding. This is in line with imaging work showing a link of the OFC/agranular insula in anxiety patients.^60^ The way to test this is to examine the role of the lOFC in associative summation in responding that is below ceiling. In our preparation, the high levels of freezing precluded from detecting summation and thus examining the role of the lOFC in this process.

Our data showing no role for the lOFC in extinction by omission is somewhat at odds with prior work showing that inactivation of lOFC disrupts extinction of fear^37^ or reward-based conditioned responding^61^. A critical difference between our work and that of others is that the overexpectation training preceded the omission part of the experiment. How learning from overexpectation and extinction by omission interact to modulate lOFC involvement is unclear. Noteworthy is the lack of agreement on the role of the lOFC in fear extinction with studies reporting disruption in extinction recall following *inactivation*^37,62^ of or NMDA-receptor *activation*^36^ in the lOFC during extinction by omission training. Although the lOFC has been linked to fear suppression in discrimination procedures^34,38,39,63^, we found no evidence of fear suppression in inactivated animals during overexpectation training nor during the start of extinction by omission training. One possibility for these discrepancies could be the anatomical location of infusion sites. While our placements spanned a very specific part of the ventrolateral OFC, those reported in other studies often extend throughout the lateral OFC and even to the adjacent agranular insular region.

As mentioned earlier, we provide additional support for the IL in fear inhibition, but we also extend the current thinking in showing that 1) the IL has a very specific role in this process; 2) the IL is not alone in supporting reduction in learned fear, and 3) the lOFC is a key player in driving fear reduction. Understanding the specific behavioural or psychological processes in the IL or the lOFC that underlie reduction in learned fear responses driven by outcome omission and overexpectation requires further investigation. But given the link between animals and humans (for review see ^60^’^64^) in the study of the brain mechanisms of fear as well as in the clinical setting, our data identify the lOFC as a potential therapeutic target for anxiety disorders (but see ^41^), and call for a rethinking of the fear circuit, thereby opening a vast novel avenue for research into the study of the basic mechanisms of fear reduction, and potentially revolutionizing the way the field currently thinks about lOFC function. Finally, taken together, our findings provide key evidence that reduction in learned fear is differentially processed in the brain even though it yields the same behavioural outcome and thus highlight the need to transcend the single-paradigm approach if a thorough understanding of the neural mechanisms of extinction is to be attained.

## Acknowledgements

This work was supported by FRQNT Nouveaux Chercheurs grant (2017-NC-198182, to MDI); a NARSAD Young Investigator grant (to MDI); a CIHR Project Grant (to MDI); the Canada Research Chairs program (to MDI); a FRQS post-doctoral fellowship (to BPPL); and a NSERC Undergraduate Student Research Award (to NB). The authors report no conflict of interest. All correspondence to be addressed to Dr. Mihaela Iordanova.

## Methods

### Subjects

One hundred and eleven (63 in Experiment 1, 48 in Experiment 2) male Sprague Dawley rats weighing between 300-370g were obtained from Harlan. Rats were pair-housed in a standard clear cage (44.5 cm × 25.8 cm × 21.7 cm) containing a mixture of beta chip and corncob bedding. The boxes were kept in an air-conditioned colony room maintained on a 12-hr light-dark cycle (lights off at 10:00 am). Food and water were available ad libitum prior to surgery and throughout the entire duration of the experiment. All experimental procedures were in accordance with the approval granted by the Canadian Council on Animal Care and the Concordia University Animal Care Committee. Rat numbers were based on other fear studies previously conducted in the lab^1^ and elsewhere.^2,3^

### Surgery and Drug Infusion

Before behavioural training and testing, rats were implanted with bilateral guide cannulae in the IL cortex or lateral OFC. Rats were anaesthetized with isoflurane gas and then mounted on a stereotaxic apparatus (David Kopf Instruments). They were then treated with a subcutaneous injection of 0.15 ml (50mg/ml) solution of rimadyl (Pfizer, Kirkland, QC) immediately upon placement in the stereotaxic frame. Twenty-two-gauge single-guide cannulae (Plastics One) were implanted through holes drilled in both hemispheres of the skull above the IL cortex or lateral OFC. The tips of the guide cannulae were aimed at the IL cortex, the following coordinates were used: 2.9 mm anterior to bregma, 2.6 mm lateral to the midline at a 30° angle (bypassing the prelimbic cortex), and 4.2 mm ventral to bregma. For the lateral OFC using the following coordinates: 3.7 mm anterior to bregma, 2.7 lateral to the midline, and 4.3 ventral to bregma. The guide cannulae were secured to the skull with four jeweller’s screws and dental cement. A dummy cannula was kept in each guide at all times except during microinjections. Rats were allowed six days to recover from surgery, during which time they were handled, weighed, and given an oral administration of 0.5 ml solution of cephalexin daily.

A cocktail of muscimol/baclofen or vehicle was infused bilaterally into the lateral OFC or IL cortex by inserting a 28-gauge injector cannula into each guide cannula. The injector cannulas were connected to a 10-μL Hamilton syringe attached to an infusion pump (Harvard Apparatus). The injector cannula projected an additional 1 mm ventral to the tip of the guide cannula. A total volume of 0.3 μl was delivered to both sides at a rate of 0.1 μl/min, and drug delivery was monitored with the progression of an air bubble in the infusion tube. The injector cannula remained in place for an additional 2 min after the infusion to allow for drug diffusion before its complete removal. Immediately after the infusion, the injector was replaced with the original dummy cannula. One day before infusions, all rats were familiarised with this procedure by removing the dummy cannula and inserting the injector cannula to minimise stress the following day.

### Drugs

A GABA_A_ agonist, muscimol (M1523, Sigma-Aldrich), and GABA_B_ agonist, baclofen (B5399, Sigma-Aldrich), were used to pharmacologically inactivate the IL cortex. A muscimol-baclofen (M/B) cocktail was prepared by dissolving 5 mg of muscimol and 93.65 mg of baclofen in 438 ml of nonpyrogenic saline (0.9% w/v) to obtain a final stock concentration of 0.1 mM muscimol-1 mM baclofen. Saline was used as a vehicle solution.

### Histology

Subsequent to behavioural testing, rats received a lethal dose of sodium pentobarbital diluted 1:1 with 0.9% sodium chloride (120 mg/kg). The brains were removed and sectioned coronally at 40 μm through the IL cortex or lateral OFC. Every second section was collected on a slide and stained with cresyl violet. The location of the cannulation tips was determined under a microscope using the boundaries defined by the atlas of Paxinos and Watson.^4^

### Behavioural Apparatus

#### Experimental chambers

Behavioural procedures were conducted in 8 operant-training chambers, each measuring 31.8 cm in height × 26.7 cm in length × 25.4 cm in width (Med Associates, St. Albans, VT, USA). The modular left and right walls were made of aluminium, and the back wall, front door, and ceiling were made of clear Perspex. Their floors consisted of stainless-steel rods, 4 mm in diameter, spaced 15 mm apart, center to center, with a tray below the floor. The grid floor was connected to a shock generator and delivered continuous scrambled foot-shock. Each chamber was enclosed in a ventilated sound attenuating cabinet. The back wall of each cabinet was equipped with a camera connected to a monitor located in another room of the laboratory where the behaviour of each rat was videotaped and observed by an experimenter. Illumination of each chamber was provided by a near-infrared light source (NIR-200) mounted on the back wall of each cabinet. Stimuli were presented through Med Associates software on a computer located outside the experimental room. The chambers had checkered or spotted wallpaper on the door and each wall with the exception of the back wall to allow for video viewing. Instead, the back wall of the holding cabinet was covered in either checkered or spotted wallpaper. The chambers (walls, ceiling, door, grid floor, and tray) were cleaned with 4% almond-scented solution (PC Black Label) after the removal of each rat. The chamber wallpaper was counterbalanced across groups.

#### Stimuli

Visual and auditory cues were used in the experiments. The visual cues consisted of a 4-Hz flashing light located on the left-hand side of the right wall and a steady light located on the right-hand side of the right wall. The auditory cues were a 70-dB tone and a 72-dB white-noise (measured inside the chamber) delivered through a loud speaker located outside the behavioural chamber. The background noise in the chamber was 48-50 dB. The background noise and experimental auditory cues were measured using a digital sound level meter (Tenma, 72-942). All cues were 30 s in duration and were fully counterbalanced. During Phases 1, 2, and 3, the cues terminated with the onset of a 1 s duration foot-shock at 0.5 mA intensity that was delivered to the floor of each chamber. The cues were controlled via a Med-Associates program.

### Behavioural Procedures

Each study was done in two replications. Allocation to groups was based on Conditioning (Phase 1 for the Overexpectation part of the study, Phase 3 for the Extinction part of the study) to ensure that all groups conditioned similarly and entered the critical part of training (Overexpectation and Extinction) with the same level of responding.

#### Phase 1 Conditioning

On days 1 to 6, rats were placed into the conditioning context for 20 min sessions. Following a 9 min 30 s adaption period, all rats received one paired presentation of the cue terminating with a foot-shock (tone on days 1, 3 and 5; flashing light on days 2, 4, and 6). On days 4 to 6, three hours following conditioning, rats were placed back in the conditioning context for 20 min where no cues were presented. These context extinction sessions were carried out in order to reduce any fear to the background cues and thus allow for a clearer assessment of the acquisition of freezing to the cues.

#### Phase 2 Overexpectation Training

Following Phase 1 Conditioning, rats were reassigned to either an overexpectation or control condition based on their responding during Conditioning. Thirty minutes prior to the start of behavioural training in Phase 2, rats received pre-training infusion of M/B or vehicle into the IL cortex (Experiment 1) or lateral OFC (Experiment 2). Following infusions, rats were further assigned to either an overexpectation or control condition, which yielded four sub conditions: overexpectation-M/B, overexpectation-vehicle, control-M/B, and control-vehicle.

On days 7, rats in the overexpectation condition were placed in the conditioning context and received two compound presentations of the tone and flashing light terminating with the onset of a foot-shock (0.5 mA, 1 s). The first trial began 4 min 30 s after being placed in the context and intertrial intervals (ITI) were 4 min 30 s. Rats remained in the chamber for 5 min following the final compound presentation. Rats in the control condition were handled for 30 s in their home-cage.

Test for *Overexpectation*. On day 8, rats were tested for responding to either the visual flashing light or auditory tone (counterbalanced). All rats were placed in the conditioning context, and after a 5 min adaptation period, the cue was presented. The test session consisted of eight stimulus alone presentations with an ITI of 1 min.

#### Phase 3 Conditioning

On day 9, rats were placed in the conditioning context, and after a 5 min adaption period, received four, 30 s paired presentations of a novel cue (steady light or white-noise, counterbalanced across all rats) and foot-shock (0.5 mA, 1 s). The ITI between paired CS-shock presentations was 3 min. Rats remained in the chamber for 2 min following the final stimulus presentation.

#### Phase 4 Extinction by omission

Following Phase 3 Conditioning, rats were reassigned to either an extinction or control condition based on their responding during Conditioning. Rats were then further assigned to either the drug or vehicle condition such that half the rats that had previously received an infusion of the drug now received vehicle, whereas the other half received drug, and similarly for rats that were initially allocated to the vehicle condition. This yielded four sub-conditions: extinction-M/B, extinction-vehicle, control-M/B, and control-vehicle. Infusions of M/B or vehicle into IL cortex (Experiment 1) or the lateral OFC (Experiment 2) occurred 30 min prior to the start of the extinction session.

On day 10, rats in the extinction condition were placed in the conditioning context, and after a 5 min adaption period, the cue was presented. The extinction session consisted of twelve, 30 s stimulus alone presentations with an ITI of 2 min (see ^5^). Rats remained in the chamber for 2 min following the final stimulus presentation. Rats in the control condition were handled for 30 s in their home-cage.

#### Test for Extinction by omission

On day 11, all rats were tested drug-free for responding to the extinguished stimulus in a manner identical to that of the extinction session.

### Data analysis

Freezing was used to assess conditioned fear. It was defined as the absence of all movements except those related to breathing.^6,7^ Each rat was observed every 2 s and scored as either freezing or not freezing by two observers, one of whom was blind to group assignment. A percentage score was calculated for the proportion of the total observation each rat spent freezing during the total duration of each CS presentation. The levels of freezing do not include freezing during the ITI. Data were analyzed in SPSS 25.0 (IBM, New York, USA) using repeated measures analysis of variance (ANOVA). Significance was set at α = 0.05. Where appropriate, adjustments with Bonferroni were made for multiple comparisons. Effects of trial, where reported, were measured with contrasts testing for the presence of a linear trend. Standardized confidence intervals (CIs; 95% for the mean difference) and measures of effect size (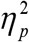 for ANOVA and Cohen’s *d* for contrasts; see Cohen^8^) are reported for each significant comparison. Data for each experiment and phase were reported using all trials. Significant outliers were detected using the Grubbs outlier test (https://www.graphpad.com/quickcalcs/Grubbs1.cfm.).

## Results

### Experiment 1: the role of the IL cortex in extinction by overexpectation and extinction by omission

#### Histology

Figures 1A and 1B shows the approximate location of injection cannulae tips for drug- and vehicle-treated rats in the overexpectation and control conditions. The plotted points represent the ventral point of the injector cannula track for each rat. Fourteen rats were excluded from statistical analysis because of incorrect placement or infection. These rats were used as anatomical controls and infusions of M/B outside of the IL cortex had no effect on the overexpectation effect (*F*_(1, 10)_ = 0.63, *p* = 0.45, 95% CI [−1.79, 1.01]) or extinction effect (*F*_(1, 7)_ = 2.41, *p* = 0.16, 95% CI [−2.35, 0.79]). This yielded the following group sizes: overexpectation-M/B, *n* = 13, overexpectation-vehicle, *n* = 11, control-M/B, *n* = 11, and control-vehicle, *n* = 14; extinction-M/B, *n* = 11, extinction-vehicle, *n* = 11, control-M/B, *n* = 8, and control-vehicle, *n* = 7.

#### Behaviour

##### Phase 1 Conditioning

The behavioural design is depicted on Figure 1C. The level of acquisition of conditioned fear, measured by freezing, to the tone and flashing light across the 6 days of conditioning was similar for each of the four groups (Figure 1D). A mixed ANOVA revealed no effect of condition (tone: *F*_(1, 45)_ = 0.18, *p* = 0.68, 95% CI [−0.55, 0.39]; flash: *F*_(1, 45)_ = 0.09, *p* = 0.77, 95% CI [−0.41, 0.52]), treatment (tone: *F*_(1, 45)_ = 0.035, *p* = 0.85, 95% CI [−0.51, 0.43]; flash: *F*_(1, 45)_ = 0.46, *p* = 0.50, 95% CI [−0.34, 0.59]), or condition x treatment interaction (tone: *F*_(1, 45)_ = 0.84, *p* = 0.36, 95% CI [−0.30, 0.64]; flash: *F*_(1, 45)_ = 0.035, *p* = 0.85, 95% CI [−0.50, 0.43]). There was a significant linear trend indicating an increase in freezing levels to the stimuli across days during conditioning (tone: *F*_(1, 45)_ = 69.77, *p* < 0.01, 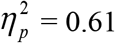, 95% CI [1.20, 1.96]; flash: *F*_(1, 45)_ = 166.96, *p* < 0.01, 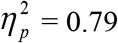, 95% CI [1.79, 2.46]), and no linear trend x condition x treatment interaction (tone: *F*_(1, 45)_ = 0.069, *p* = 0.79, 95% CI [−0.84, 1.04]; flash: *F*_(1, 45)_ = 0.041, *p* = 0.84, 95% CI [−0.75, 0.88]). These results indicate the overall levels of freezing and rate of acquisition was similar across groups.

##### Phase 2 Overexpectation Training

Figure 1E shows responding to the compound presentations during overexpectation training for the rats in the overexpectation condition. Infusions of M/B in the IL cortex had no effect on within-session responding to the compound stimulus. A mixed ANOVA revealed no main effect of treatment (*F*_(1, 22)_ = 0.022, *p* = 0.88, 95% CI [−0.64, 0.74]), a significant linear trend (*F*_(1, 22)_ = 15.10, *p* = 0.001, 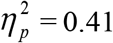, 95% CI [0.44, 1.44]) and no linear trend x treatment interaction (*F*_(1, 22)_ = 1.45, *p* = 0.24, 95% CI [−1.59, 0.42]). The data confirm that rats in the overexpectation-M/B and overexpectation-vehicle maintained similar responding to the compound stimulus during this phase.

##### Test for Overexpectation

Figure 1F shows the levels of freezing to the target stimulus during Test for each of the four groups. There was no effect of IL inactivation in overexpectation. A mixed ANOVA revealed a main effect of condition (*F*_(1, 45)_ = 19.21, *p* < 0.001, 95% CI [−1.48, −0.41], d = 1.30), no main effect of treatment (*F*_(1, 45)_ = 0.33, *p* = 0.57, 95% CI [−0.66, 0.41]), and no condition x treatment interaction (*F*_(1, 45)_ = 0.21, *p* = 0.65, 95% CI [−0.64, 0.44]). There was a significant linear trend across trials (*F*_(1, 45)_ = 42.63, *p* < 0.001, 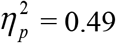, 95% CI [−1.01, −0.53]), confirming a decline in responding across Test but there was no linear trend x condition x treatment interaction (*F*_(1, 45)_ = 0.24, *p* = 0.63, 95% CI [−0.47, 0.70]), confirming a similar decline in responding across all groups. These results indicate that infusion of M/B in the IL cortex prior to compound presentations during overexpectation training had no effect on retrieval of the overexpectation memory when tested drug-free.

##### Phase 3 Conditioning

The behavioural design is depicted on Figure 2C. The rate and level of acquisition was consistent across all groups (Figure 2D). A mixed ANOVA revealed no main effect of condition (*F*_(1, 33)_ = 0.19, *p =* 0.67, 95% CI [−0.52, 0.74]), treatment (*F*_(1, 33)_ = 0.06, *p* = 0.81, 95% CI [−0.69, 0.57]), nor condition x treatment interaction (*F*_(1, 33)_ = 0.038, *p* = 0.85, 95% CI [−0.58, 0.68]). There was a significant linear trend indicating an increase in conditioned freezing response across conditioning (*F*_(1, 33)_ = 44.82, *p* < 0.01, 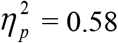, 95% CI [0.76, 1.42]), and no linear trend x condition x treatment interaction (*F*_(1, 33)_ = 0.84, *p* = 0.37, 95% CI [−1.12, 0.52]). These results indicate that the overall levels and rate of acquisition was similar across groups.

##### Phase 4 Extinction by omission

Figure 2E shows the levels of freezing to the cue undergoing extinction under vehicle versus M/B inactivation of the IL cortex. Overall, rats infused with M/B in the IL cortex prior to extinction training froze significantly more to the cue compared to vehicle-treated rats (*F*_(1, 20)_ = 12.80, *p* = 0.002, 95% CI [0.40, 1.54], d = 1.53). These results indicate that silencing the IL cortex disrupted the loss in conditioned responding across extinction training. There was a significant linear trend across trials (*F*_(1, 20)_ = 9.49, *p* = 0.006, 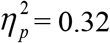, 95% CI [−1.18, −0.23]) but no linear trend x treatment interaction (*F*_(1, 20)_ = 0.67, *p* = 0.42, 95% CI [−1.33, 0.58]), indicating that the rate of reduction in responding was similar for both extinction groups.

##### Test for Extinction by omission

Figure 2F shows the levels of freezing to the cue on Test following extinction training. Inactivation of the IL cortex prior to extinction training disrupted retrieval of the extinction memory on Test. A mixed ANOVA revealed no main effect of condition (*F*_(1, 33)_ = 2.23, *p* = 0.15, 95% CI [−0.84, 0.25]), a main effect of treatment (*F*_(1, 33)_ = 4.52, *p* = 0.041, 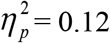, 95% CI [−0.12, 0.92], d = 0.89), and a significant condition x treatment interaction (*F*_(1, 33)_ = 4.72, *p* = 0.037, 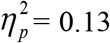, 95% CI [−0.11, 0.97]). Post hoc tests with Bonferroni adjustments revealed a significant difference within the extinction conditions: rats treated with M/B during extinction froze significantly more during presentations of the CS at Test compared to rats treated with vehicle (*F*_(1, 33)_ = 11.43, *p* = 0.002, 95% CI [0.16, 1.54], d = 1.85); there was no effect of drug on the control condition (*F*_(1, 33)_ = 0.001, *p* = 0.98, 95% CI [−0.85, 0.83]). There was a significant linear trend across trials (*F*_(1, 33)_ = 40.05, *p* < 0.001, = 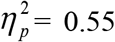, 95% CI [−1.24, −0.64]) but no linear trend x condition x treatment interaction (*F*_(1, 33)_ = 0.07, *p =* 0.79, 95% CI [−0.89, 0.73]). These results indicate that infusion of M/B in the IL cortex prior to extinction training disrupted retrieval of the extinction memory on in the drug-free Test.

### Experiment 2: the role of the OFC in extinction by overexpectation and extinction by omission

#### Histology

Figure 3A and 3B shows the approximate location of injection cannulae tips for drug- and vehicle-treated rats in the overexpectation and control conditions. The plotted points represent the ventral point of the injector cannula track for each rat. Three rats were excluded from statistical analysis because of incorrect placement or infection. Moreover, one rat was identified as a significant outlier on Test for Overexpectation using the Grubb’s outlier test (p < 0.05) and thus excluded from all statistical analyses. This yielded the following group sizes: overexpectation-M/B, *n* = 12, overexpectation-vehicle, *n* = 11, control-M/B, *n* = 12, and control-vehicle, *n* = 9; extinction-M/B, *n* = 10, extinction-vehicle, *n* = 12, control-M/B, *n* = 11, and control-vehicle, *n* = 11.

#### Behaviour

##### Phase 1 Conditioning

The behavioural design is depicted on Figure 3C. The level of acquisition of conditioned fear, measured by freezing, to the tone and flashing light across the 6 days of conditioning was similar for each of the four groups (Figure 3D). A mixed ANOVA revealed no effect of condition (tone: *F*_(1, 40)_ = 0.001, *p* = 0.97, 95% CI [−0.46, 0.48]; flash: *F*_(1, 40)_ = 0.52, *p* = 0.48, 95% CI [−0.39, 0.70]), treatment (tone: *F*_(1, 40)_ = 0.53, *p* = 0.47, 95% CI [−0.33, 0.60]; flash: *F*_(1, 40)_ = 0.76, *p* = 0.39, 95% CI [−0.35, 0.73]), or condition x treatment interaction (tone: *F*_(1, 40)_ < 0.001, *p* = 0.98, 95% CI [−0.46, 0.47]; flash: *F*_(1, 40)_ = 0.66, *p* = 0.42, 95% CI [−0.37, 0.72]). There was a significant linear trend indicating an increase in freezing levels to the stimuli across days during conditioning (tone: *F*_(1, 40)_ = 213.40, *p* < 0.01, = 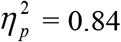, 95% CI [2.51, 3.31]; flash: *F*_(1, 40)_ = 264.90, *p* < 0.01, 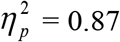, 95% CI [2.73, 3.50]), and no linear trend x condition x treatment interaction (tone: *F*_(1, 40)_ = 0.15, *p* = 0.70, 95% CI [−0.84, 1.15]; flash: *F*_(1, 40)_ = 0.007, *p* = 0.93, 95% CI [−0.92, 0.99]). These results indicate the overall levels of freezing and rate of acquisition was similar across groups.

##### Phase 2 Overexpectation Training

Figure 3E shows responding to the compound presentations during overexpectation training for the rats in the overexpectation condition. Infusions of M/B in the lateral OFC had no effect on within-session responding to the compound stimulus. A mixed ANOVA revealed no main effect of treatment (*F*_(1, 21)_ = 0.05, *p* = 0.83, 95% CI [−0.60, 0.75]), no linear trend (*F*_(1, 21)_ = 1.42, *p* = 0.25, 95% CI [−0.23, 0.86]) and no linear trend x treatment interaction (*F*_(1, 21)_ = 2.40, *p* = 0.14, 95% CI [−1.90, 0.28]). The data confirm that rats in the overexpectation-M/B and overexpectation-vehicle maintained similar responding to the compound stimulus during this phase.

##### Test for Overexpectation

Figure 3F shows the levels of freezing to the target stimulus during Test for each of the four groups. Inactivation of the lateral OFC prior to overexpectation training disrupted the overexpectation effect on Test. A mixed ANOVA revealed no main effect of condition (*F*_(1, 40)_ = 0.42, *p* = 0.52, 95% CI [−0.73, 0.45]) or treatment (*F*_(1, 40)_ = 2.53, *p* = 0.12, 95% CI [−0.24, 0.94]), but there was a significant condition x treatment interaction (*F*_(1, 40)_ = 8.73, *p* = 0.005, = 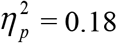, 95% CI [0.054, 1.23]). Post hoc tests with Bonferroni adjustments revealed a significant difference within the overexpectation conditions: rats treated with M/B froze significantly more during presentations of the target stimulus at Test compared to vehicle-treated rats (*F*_(1, 40)_ = 10.93, *p* = 0.002, 95% CI [0.18, 1.80], d = 1.44); there was no effect of drug on the control condition (*F*_(1, 40)_ = 0.88, *p* = 0.35, 95% CI [−1.15, 0.56]). There was a significant linear trend across trials (*F*_(1, 40)_ = 84.59, *p* < 0.001, 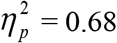, 95% CI [−1.57, −1.00]), confirming a decline in responding across Test but there was no linear trend x condition x treatment interaction (*F*_(1, 40)_ = 2.84, *p* = 0.10, 95% CI [−1.23, 0.29]), confirming a similar decline in responding across all groups. These results indicate that infusion of M/B in the lateral OFC prior to compound presentations during overexpectation training disrupted retrieval of the overexpectation memory when tested drug-free.

##### Phase 3 Conditioning

The behavioural design is depicted on Figure 4C. Acquisition of fear to the cue was equivalent between all groups (Figure 4D). A mixed ANOVA revealed no main effect of condition (*F*_(1, 40)_ = 0.003, *p =* 0. 96, 95% CI [−0.57, 0.55]), treatment (*F*_(1, 40)_ = 0.16, *p* = 0.69, 95% CI [−0.65, 0.47]), or condition x treatment interaction (*F*_(1, 40)_ = 0.015, *p* = 0.90, 95% CI [−0.59, 0.54]). There was a significant linear trend indicating an increase in conditioned freezing response across conditioning (*F*_(1, 40)_ = 67.52, *p* < 0.01, 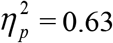, 95% CI [0.93, 1.53]), and no linear trend x condition x treatment interaction (*F*_(1, 40)_ = 0.001, *p* = 0.98, 95% CI [−0.76, 0.74]). These results indicate that the overall levels and rate of acquisition was similar across groups.

##### Phase 4 Extinction by omission

Figure 4E shows the levels of freezing to the cue undergoing extinction, i.e., non-reinforced presentations, under vehicle versus M/B inactivation of the lateral OFC. Infusion of M/B in the lateral OFC prior to extinction training had no effect on within-session performance. Levels of freezing to the cue across extinction did not differ between extinction-M/B and extinction-vehicle rats (*F*_(1, 20)_ = 0.90, *p* = 0.35, 95% CI [−0.85, 0.32]). There was a significant linear trend across trials (*F*_(1, 20)_ = 35.80, *p* < 0.001, 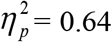, 95% CI [−1.55, −0.75]) but no linear trend x treatment interaction (*F*_(1, 20)_ = 2.33, *p* = 0.14, 95% CI [−0.22, 1.38]), indicating that the rate of reduction in responding was similar for both extinction groups.

##### Test for Extinction by omission

Figure 4F shows the levels of freezing to the cue on Test following extinction training. Overall, extinction training reduced fear on Test, but this extinction effect was not modulated by inactivation of the lateral OFC during extinction learning. A mixed ANOVA revealed a main effect of condition (*F*_(1, 40)_ = 13.65, *p* = 0.001, 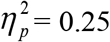, 95% CI [−0.88, −0.14], d = 1.12) but no main effect of treatment (*F*_(1, 40)_ = 0.79, *p* = 0.38, 95% CI [−0.25, 0.50]), and no condition x treatment interaction (*F*_(1, 40)_ = 0.90, *p* = 0.35, 95% CI [−0.50, 0.24]). There was a significant linear trend indicating a reduction in conditioned freezing to the cue on Test across trials (*F*_(1, 40)_ = 79.81, *p* < 0.001, 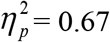, 95% CI [−1.53, −0.96]) but no linear trend x condition x trial interaction (*F*_(1, 40)_ =, *p =* 0.67, 95% CI [−0.63, 0.88]), confirming a similar decrease in responding across trials for all groups. These results indicate that inactivating the lateral OFC with M/B prior to extinction training had no effect on extinction learning.

